# Effect of chemical modifications of tannins on their antibiofilm effect against Gram-negative and Gram-positive bacteria

**DOI:** 10.1101/2022.05.26.493672

**Authors:** Xabier Villanueva, Lili Zhen, José Nunez Ares, Thijs Vackier, Heiko Lange, Claudia Crestini, Hans Steenackers

**Author notes:** Affiliated with 2) via NAST – Nanoscience & Nanotechnology & Innovative Instrumentation Center.

## Abstract

**Background:** Tannins have demonstrated antibacterial and antibiofilm activity, but the mechanisms of action are not completely elucidated. We are interested in understanding how to modulate the antibiofilm activity of tannins and in delineating the relationship between chemical determinants and antibiofilm activity.

**Materials and methods:** the effect of five different naturally acquired tannins and their chemical derivatives on biofilm formation and planktonic growth of *Salmonella* Typhimurium, *Pseudomonas aeruginosa, Escherichia coli* and *Staphylococcus aureus* was determined in the Calgary biofilm device.

**Results:** most of the unmodified tannins exhibited specific antibiofilm activity against the assayed bacteria. The chemical modifications were found to alter the antibiofilm activity level and spectrum of the tannins, with the positive charge introducing C_3_NMe_3_Cl-0.5 derivatization shifting the anti-biofilm spectrum towards Gram-negative bacteria and C_3_NMe_3_Cl-0.1 and the acidifying CH_3_COOH derivatization shifting the spectrum towards Gram-positive bacteria. Also, the quantity of phenolic-OH groups per molecule has a weak impact on the anti-biofilm activity of the tannins.

**Conclusions:** we were able to modulate the antibiofilm activity of several tannins by specific chemical modifications, providing a first approach for fine tuning of their activity and spectrum.

## Introduction

Plant-derived tannins have been used from ancient times in leather industry because of their ability of making leather last for a long time, and the name “tannin” comes from the use of these chemicals for “tanning” leather (1, 2). Accordingly, it was hypothesized that the resistance of leather to microbial decomposition could be explained by this use of tannin-rich compounds in leather curation and several studies have pointed to multiple additional pharmacological properties of tannins, including anti-inflammatory and anti-cancer effects, which are contributing to a renewed interest in these products as a source of new bio-based pharmaceuticals (1, 3–5). Tannins have been shown to inhibit bacterial growth of different Gram-positive and Gram-negative bacteria (4, 6–9), and are shown to be able to disperse biofilms (10). On some occasions this antibiofilm activity is specific and independent from the ability to inhibit bacterial growth (11). Examples of tannins with antibacterial activity are tannic acid (12), ellagic acid (13) and epigallocatechin gallate (14).

From a molecular point of view, tannins can be divided in two groups: condensed and hydrolysable tannins. Hydrolysable tannins are esters of gallic acid with a core sugar, often glucose or quinic acid. Tannic acid is the most prominent representative of the family of hydrolysable tannins comprising a glucose center (15, 16). Condensed tannins are oligomeric and polymeric proanthocyanidins, consisting of flavan-3-ol units, linked by carbon–carbon bonds not susceptible to hydrolytic cleavage (16, 17). The scaffold of the subclass of tannins called complex tannins is very similar to those found in condensed and hydrolysable tannins, where a flavan-3-ol unit is linked to gallic acid in a monomeric or polymeric system (2). However, there is no reference in literature that this kind of differentiation has any effect on the level and kind of antimicrobial activity of the tannins.

To date, there is no clear understanding of the antimicrobial mechanisms of action of the different tannins. One early hypothesis suggested that the ability of tannins to form complexes with leather proteins is underlying their mechanism of antimicrobial action (18). It has been suggested that the observable activity could be explained by the presence of free phenolic hydroxyl groups which can affect, for example, enzymatic activity via covalent or non-covalent linking (19). In this respect, it has been seen that phenolic compounds can have antimicrobial effects against *Pseudomonas aeruginosa* and *Staphylococcus aureus* (20, 21). This ultimately means that the typical phenolic character of the tannins could play an important role for the antimicrobial activity (22, 23). Other mechanisms of action for the antimicrobial activity of tannins have also been described, in particular for tannic acid, like disruption of peptidoglycan formation (24), iron chelation (12), membrane disruption (25), efflux pump inhibition (26) and fatty acid synthesis (27).

It has also been shown in previous literature that tannins have the ability to reduce biofilm formation (20, 21, 28, 29). Biofilms are conglomerates of bacteria, usually at an interphase (solid-air, solid-liquid, liquid-air), that are surrounded by a protective mesh of extracellular polymeric substances (EPS). This enhances the ability of the bacteria to survive dehydration, disinfectants and antibiotics (30, 31). The mechanisms of protection include reduced penetration of antimicrobial compounds, reduced bacterial metabolism, induction of efflux pumps and more frequent horizontal gene transfer (31).

Regarding the antibiofilm activity of tannins, it has been described that some of them have a biofilm-specific mechanism of action, such as inhibition of quorum sensing (QS) in *P. aeruginosa* by the tannin-rich fractions of *Terminalia catappa* (32) and *T. chebulata* (33), and induction of transglucosylase activity in *S. aureus* by tannic acid (34). This type of biofilm-specific behavior is actually desired, because the lack of direct growth inhibition decreases the selective pressure towards resistance phenotypes (35– 37).

Because of this reduced potential of resistance development, in the current study we evaluated selected chemical variants of tannins for their ability to inhibit biofilm formation without inhibiting planktonic growth. Also, we investigated how different chemical modifications change the activity level and spectrum of the tannins. Understanding the effect of structural features on activity allows to enhance or finetune activity. The scarcity of structure-activity relationship (SAR) research in the field of tannins and bioactive phenolic compounds from plant sources as antimicrobial compounds highlights the value of this work (38, 39).

## Materials and Methods

### Assayed tannins

Five commercially available tannin extracts, comprising three condensed and two hydrolysable tannins were used. The three condensed tannins comprised Omnivin 20R (monomeric (epi)catechin, **Vv- 20**), Omnivin WG (procyanidins (62%)/profisetidins (34%), **Vv**) and Mimosa ATO ME (prorobinetidins (33%)/profisetidins (67%), **Am**), and the two hydrolysable tannins Tanal 01 (tannic acid, **Ta-01**) and Tanal 04 (galloylquinic acid, **Ta-04**). The chemical structures of these tannins were elucidated in detail as reported elsewhere (40). Fig. 1A gives an overview of the structural features. As can be seen in the figure, the tannins **Vv-20** and **Vv** comprise low molecular size monomeric or oligomeric tannins, and the tannins **Am, Ta-01** and **Ta-04** are polymeric tannins of larger molecular size. The five selected tannins were chemically modified by derivatizing them via their phenolic functionalities, as reported in detail elsewhere (41), with different levels of specific functional motifs: i) hydroxy-*N,N,N*-trimethylpropanyl-3- aminium chloride (**C**_**3**_**NMe**_**3**_**Cl-eq**), ii) hydroxypropyl-1-carboxylic acid (**C**_**3**_**COOH-eq**), and iii) oligomeric ethylene glycol polyether (**PEG**_**500**_**-eq**), whereby ‘eq.’ in the compounds listed in Fig. 1B indicates the equivalents of the functional motif that were used for the chemical modification (41). This chemical functionalizations gave several properties to the tannins: i) **C**_**3**_**NMe**_**3**_**Cl-eq** added positive charges to the tannin molecule, ii) **C**_**3**_**COOH-eq**, as a weak acid, potentially added negative charges to the molecule, and iii) **PEG**_**500**_**-eq** polymerized the tannin molecules. During the various functionalizations, control tannins were re-isolated from blank reactions. Tannins labelled as ‘**Blank-W**’ are tannins isolated from blank reactions performed in water and tannins labelled as ‘**Blank-D**’ are tannins isolated from blank reaction preformed in dimethylformamide.

**FIG 1.**
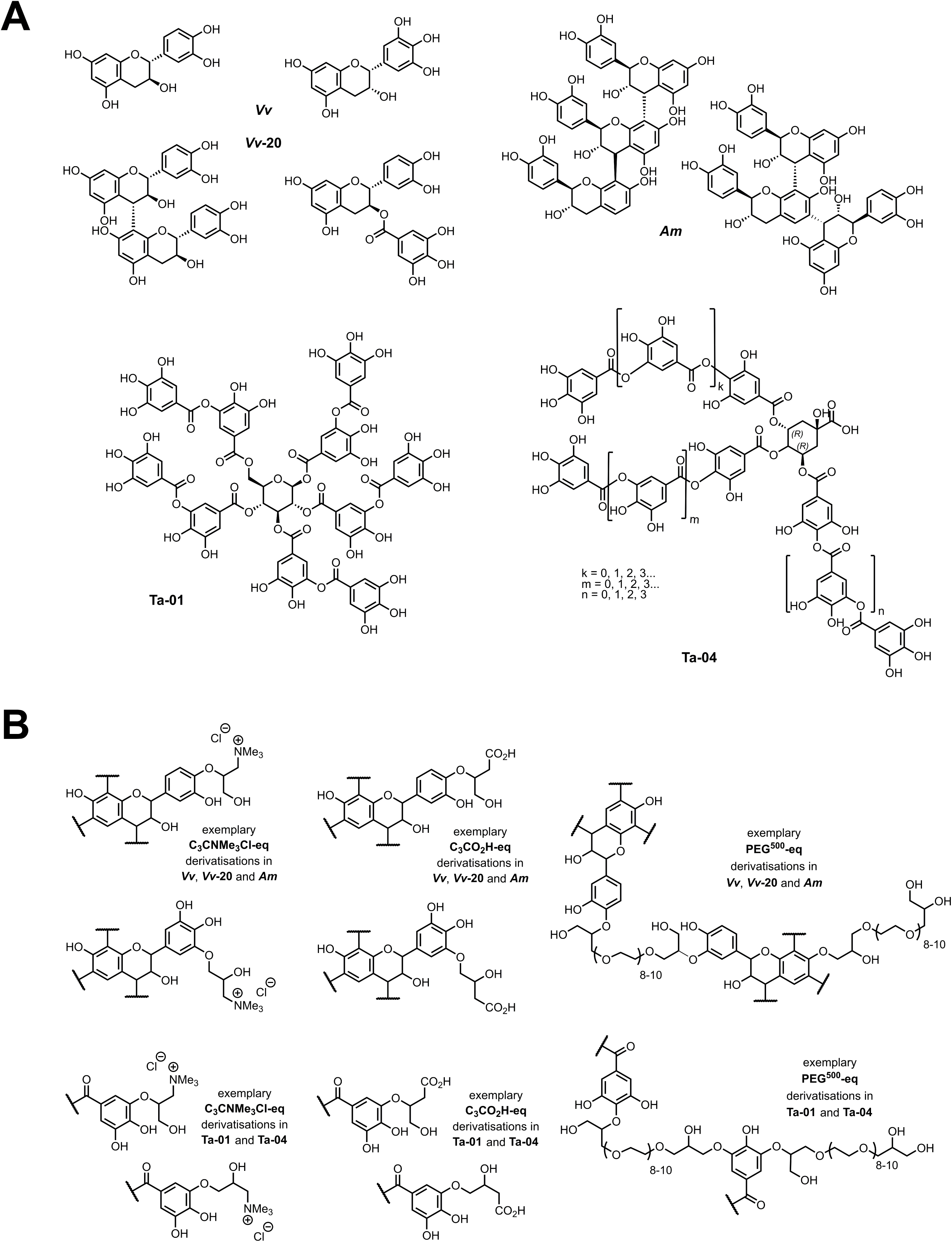
(A) Chemical structure of commercially available condensed and hydrolysable tannins used in this study (40) and (B) their chemically derivatized structures as described elsewhere (41). Exemplary structural aspects are shown; synthetic route leads to generation of both primary and secondary aliphatic alcohols within the total C_3_ linker moiety connecting the functional to the tannin (41). Legend: **C**_**3**_**NMe**_**3**_**Cl-eq** - hydroxy-*N,N,N*-trimethylpropanyl-3-aminium chloride; ii) **C**_**3**_**COOH-eq** - hydroxypropyl-1- carboxylic acid, and iii) **PEG**_**500**_**-eq** - oligomeric ethylene glycol polyether (PEG_500_-eq).

Because of the low solubility of the tannins in aqueous media, the dry compounds were first suspended in dimethyl sulfoxide (DMSO) at a stock concentration of 60 g/l and from there diluted to the desired concentration in the following experiments. A 1% v/v concentration of DMSO was never exceeded in order to prevent potential effects of DMSO on bacterial growth or biofilm formation. All the tests were done in aerobic conditions in growth media with a pH of ∼ 7.4 and a salinity range of 0.025-0.5% w/v.

### Antibiofilm assay

Biofilms of *Salmonella enterica* var. Typhimurium ATCC14028, *Pseudomonas aeruginosa* PA14, *Escherichia coli* TG1 and *Staphylococcus aureus* SH1000 were grown in the Calgary biofilm device via a protocol that was previously described (42, 43). The bacteria were grown overnight (ON) in LB broth at 37°C. These ON cultures were then diluted 1/100 in diluted (1/20) Tryptic Soy Broth (TSB, Thermo Fisher Scientific) for Gram-negative bacteria and in undiluted TSB for *S. aureus*. 100 µL of growth medium or a solution of the tannins in growth medium was added to the wells of a 96-well Calgary device. The diluted ON cultures were then added to the wells to obtain a starting inoculum of 10^6^ cfu/ml in a final volume of 200 µl of growth medium per well. Also, both for exploratory screening test and validation of anti-biofilm activity, one row of the plate was filled with inoculum with growth media without tannin and another row was filled with media without bacteria. To account for potential effects of the compounds themselves, the same tannin-concentration was added to separate control 96-well Calgary biofilm plates without the inoculation of the bacteria. Afterwards, all plates were incubated in a wet chamber for the appropriate time and temperature: 48 h at 37 °C for *S. aureus*, 24 h at 25°C for the Gram-negatives.

Biofilm formation and planktonic growth were determined by crystal violet staining of the pegs and OD_600_ measurements of the base plate of the device, respectively. Specifically, after incubation the covers of the plates (which contain the pegs) were removed and washed once with PBS. The pegs were then stained with 200 µl per well of 0.1% v/v of crystal violet (CV, VWR International) for 30 minutes. After staining, the excess of CV was washed once with 200 µl per well of distilled water and let dry for 30 minutes. Finally, the CV was recovered in a new 96-well plate with 200 µl per well of 30% v/v glacial acetic and the optical density at 570 nm (OD_570_) measured in a plate reader. The optical density at 600 nm (OD_600_) of the bacteria in the base plates was measured to determine the growth of the planktonic cells.

### Exploratory antibiofilm screening

In a preliminary experiment, the tannins were tested in two-fold dilution series ranging from 600 to 9.38 mg/l in 3 independent replicates. The obtained raw OD_570_ (for the biofilm formation) and OD_600_ (for the planktonic growth) were corrected by the OD of the tannins incubated in absence of bacterial inoculum and then normalized using the average OD of the bacterial inoculum incubated in absence of tannin, thus being converted to percentage of biofilm formation (OD_570_) and percentage of bacterial growth (OD_600_).

The BIC_50_ and IC_50_ (the compound concentration required to inhibit the biofilm formation and the bacterial growth by 50%, respectively) were calculated by applying a log[tannin] vs percentage of biofilm formation or percentage of planktonic growth non-linear regression (four-parameters) using the statistical package GraphPad 8.0.

### Validation anti-biofilm screening: experimental design and statistical analysis

Based on the information from the preliminary experiment, a definitive antibiofilm experiment in the Calgary biofilm device was set up with 8 independent repeats. We first defined the factors under study, i.e., the parameters that potentially have an influence on the formation of biofilm. The considered parameters are: (i) Original unmodified tannin: **Vv-20, Vv, Ta-01, Ta-04, Am**, (ii) Concentration of tannin (mg/l): 9.38, 79.69, 150, (iii) Chemical modifier: **C**_**3**_**COOH** (AC), **PEG, C**_**3**_**NMe**_**3**_**Cl** (AM), **Blank-D** (blank reaction with dimethylformamide), **Blank-W** (blank reaction with water), Unmodified and (iv) Concentration of applied chemical modifier: Low and High (Low: 0.05 and 0.1 Eq of chemical substitution; High: 0.25 and 0.5 Eq of chemical substitution). In order to minimize effects of plate-to-plate variation, we applied an optimal randomized experimental design with the statistical package JMP 15.0 to reduce experimental noise and confounding factors. Different to the preliminary experiment, only three concentrations were assayed, which were chosen to best capture the antibiofilm activity of the tannins: 9.38, 79.69 and 150 mg/l. The remaining part of the protocol of this experiment is identical to that of the preliminary experiment described above.

To determine the effect of the chemical derivatizations on the antibiofilm and antibacterial effect of the unmodified tannins on each of the assayed bacteria, an ANOVA test with Tukey post-hoc test comparing the unmodified tannins with their respective derivatized tannins was done based on the obtained biofilm formation and planktonic growth levels.

### Relationship between the phenolic hydroxyl content and antibiofilm effect

To determine the effect of phenolic hydroxyl (OH) content on the antimicrobial and antibiofilm effect of the different tannins, a simple linear regression between the phenolic OH content of the tannins and the level of biofilm formation (or planktonic growth) was performed. To better calculate this correlation, we used the different tannin assayed concentrations and the obtained mmol of phenolic OH per gram of material to calculate the mmol of phenolic OH present in the system for each tannin at each assayed concentration, and we correlated this value with the respective percentage of biofilm inhibition and planktonic growth inhibition. The phenolic OH content of the tannins was determined via ^31P^NMR as described elsewhere (40).

## Results and discussion

### Exploratory screening to determine the concentration test range

In order to delineate a structure-activity and functionality-activity relationship, a diverse range of five commercially obtained tannins, three condensed tannins and two hydrolysable tannins, were derivatized with different levels of three functional motifs. The condensed tannins were previously identified as mixtures of (epi)catechins and fisetinidols (41) and consisted of (i) the essentially monomeric **Vv-20**, (ii) the low oligomeric **Vv** and (iii) the higher oligomeric **Am**. The hydrolysable tannins consisted of two large tannins: (i) the “typical” tannic acid **Ta-01** and (ii) the galloquinic acid derivative **Ta-04**. All tannins were previously functionalized with a positive charge introducing ammonium salt hydroxy-*N,N,N*-trimethylpropanyl-3-aminium chloride (**C**_**3**_**NMe**_**3**_**Cl-eq**), an acidifying hydroxypropyl-1- carboxylic acid (**C**_**3**_**COOH-eq**) motive and a polymerizing oligomeric ethylene glycol polyether (**PEG**_**500**_**-eq)** (Fig. 1). The preventive activity of both the natural and derivatized tannins against the biofilm formation and planktonic growth of the Gram-negative species *S*. Typhimurium, *P. aeruginosa, E. coli* and the Gram-positive species *S. aureus* was evaluated by means of the Calgary biofilm device. In a first set of exploratory experiments, two-fold serial dilutions (from 600 till 4.96 mg/l) of the tannins were evaluated in order to obtain a first glance on activity spectrum and active concentration range. Activities against all four bacterial species were observed, with BIC_50_ values ranging from 4.69 to 545.8 mg/l and IC_50_ values ranging from 37.5 to 459.2 mg/l (Table S1 in ‘Supplementary Material’). As such this experiment allowed to determine the test concentrations for future validation experiments: 9.38 mg/l was the lowest assayed concentration in the preliminary screening and the most active tannins exhibited BIC_50_ equal to or below that value; 150 mg/l was the concentration at which almost all tannins with antibiofilm effect were active; and 79.69 mg/l is the average of those two concentrations. Such validation experiments were required because the exploratory experiments only had three independent repeats and this did not provide sufficient statistical power to distinguish the levels of activity of the different tannins and delineate the relationship between the chemistry and the antibiofilm level of the tannins. Furthermore, there was a statistically significant plate-to-plate variation between the controls of each plate (see Figures S1 and S2 in ‘Supplementary Material’).

### Extensive randomized validation screening at limited number of concentrations

To allow a multivariate analysis considering tannin scaffold, derivatization and concentration as well as bacterial target species, the previous antibiofilm and antimicrobial experiments were repeated in one experiment with eight repeats per condition, but only focusing on the three tannin concentrations that could capture best the antibiofilm effect of the tannins: 9.38, 79.69 and 150 mg/l. In order to minimize previously observed effects of plate-to-plate variation, these experiments were designed in a randomized way, i.e., all the tannins were distributed through all the plates in a random fashion. This allowed to decrease the random error and the possibility of confounding factors, a necessary step for doing a complex statistical analysis that allows to link the different chemical characteristics. In what follows we will first focus on the unmodified tannins, after which we will elaborate on the effect of chemical derivatization.

### Effect of unmodified tannin scaffold on the antibiofilm activity level and spectrum

In Fig. 2 it can be seen that the commercially available natural tannins showed different antibiofilm activities against the four assayed bacterial species. All unmodified tannins can be considered to have “broad spectrum activity”, since all of them exhibited statistically significant antibiofilm activity against Gram-positive and Gram-negative bacteria at least at the concentration of 150 mg/l, according to an ANOVA test with Tukey post-hoc analysis. Also, most of the assayed tannins exhibited a concentration-dependent activity, depending on the assayed tannin and the tested bacteria.

**FIG 2.**
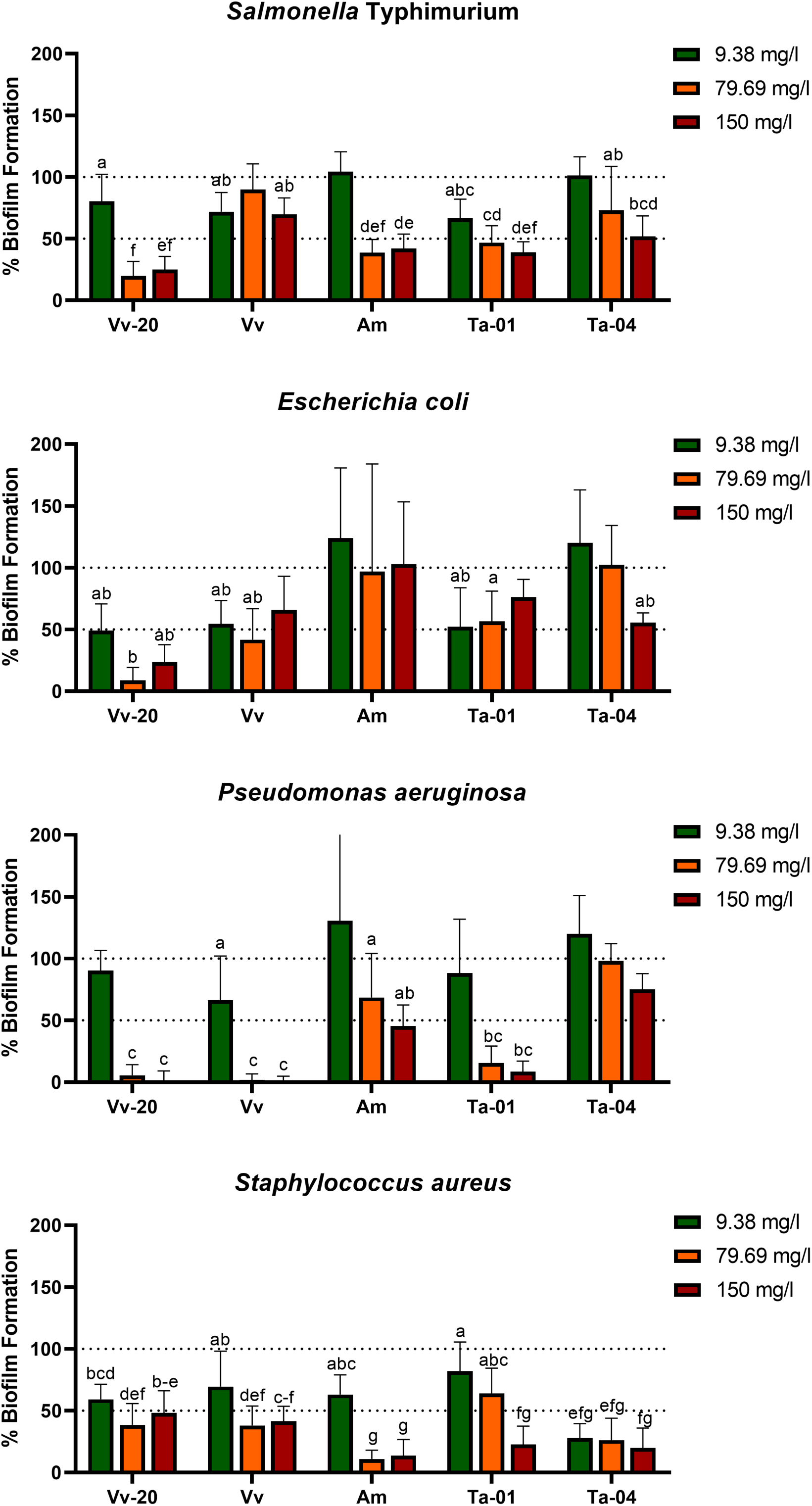
Biofilm formation (expressed as percentage in comparison to control) in the presence of 9.38, 79.69 and 150 mg/l of unmodified tannins. The letters indicate groups of tannin-concentration combinations whose effects are significantly different from the control but not significantly different to each other; the bars with no letter are those tannin-concentration combinations which are not significantly different from the control. The statistical differences were determined via ANOVA test with Tukey post-hoc analysis, with a p value of 0.05.

However, there were clear differences in the degree of antibiofilm activity of each tannin. Starting from the condensed tannins, the monomeric **Vv-20** exhibited significant antibiofilm activity against all the assayed bacteria at all the assayed concentrations in a clear dose-dependent way, a dose of 79.69 mg/L being sufficient for inhibiting biofilm formation more than 50% against the four assayed bacteria. Also, **Vv-20** was the most effective tannin against *S*. Typhimurium, with more than 75% of biofilm inhibition at 79.69 and 150 mg/l, and against *E. coli*, with more than 80% biofilm inhibition at 79.69 mg/l. Regarding the low oligomeric **Vv**, the antibiofilm activity was preferential against *P. aeruginosa* and *S. aureus*, for which it exhibited potent dose-dependent antibiofilm activity. This compound was able to inhibit more than 50% of biofilm formation by *S. aureus* both at 79.69 and 150 mg/l, and is the unmodified tannin with highest effect against *P. aeruginosa*, displaying antibiofilm activity of more than 90% at 79.69 mg/l and more than 95% at 150 mg/l. On the contrary, **Vv** was less effective against *S*. Typhimurium and *E. coli* and was able to inhibit less than 30% of biofilm formation of both bacterial species, regardless of the assayed concentration. Contrary to the previous two unmodified condensed tannins, the high oligomeric **Am** has preferential antibiofilm activity against *S*. Typhimurium and *S. aureus*, but also has significant antibiofilm activity against *P. aeruginosa*. Moreover, **Am** is the unmodified tannin with the highest antibiofilm activity against *S. aureus*, with more than 85% of antibiofilm activity at 79.69 and 150 mg/l, also inhibiting more than 50% of biofilm formation of *S*. Typhimurium both at 79.69 and 150 mg/l, and inhibiting *P. aeruginosa* biofilm formation in more than 30% at 79.69 mg/l and more than 50% at 150 mg/l. On the contrary, **Am** has unnoticeable inhibitory activity against *E. coli* biofilms at any of the assayed concentrations.

With respect to the hydrolysable tannins, tannic acid, **Ta-01** exhibited preferential activity against *P. aeruginosa* and *S*. Typhimurium. The biofilm inhibitory activity of **Ta-01** ranged from 40% at 9.38 mg/l to 60% at 150 mg/l against *S*. Typhimurium, and from 15% at 9.38 mg/l to more than 90% at 150 mg/l against *P. aeruginosa*. On the contrary, the galloquinic acid derivative, **Ta-04** exhibited preferential activity against *S. aureus*. While the antibiofilm activity of **Ta-04** against *S*. Typhimurium and *E. coli* was dose dependent (reaching a maximum of 50% biofilm inhibition against both bacteria at a concentration of 150 mg/l), the antibiofilm activity of **Ta-04** against *S. aureus* was not dose dependent, with more than 70% biofilm inhibition at the 3 assayed concentrations.

Importantly, the unmodified tannins were in general not found to have antibacterial activity against the planktonic bacteria, except for the low oligomeric condensed **Vv** against *S*. Typhimurium and the hydrolysable tannic acid **Ta-01** against *E. coli* at the highest concentration of compound (Fig. 3). In previous reports, it was found that hydrolysable tannins similar to galloylquinic acid, hence similar to **Ta- 04**, exhibit broad spectrum antibiofilm activity (3, 15, 44). Those same reports, however, also suggest that hydrolysable tannins have antibacterial effects against planktonic Gram positive and Gram bacteria, which was not observed in our experiment. On the other hand, tannic acid (45) and 1,2,3,4,6-penta-*O*- galloyl-β-D-glucopyranose (46), both hydrolysable tannins, have been reported to inhibit biofilm formation of *S. aureus* without inhibiting planktonic growth, which is consistent with the results observed for tannic acid **Ta-01**. In our assay, also galloylquinic acid **Ta-04** showed such activity. This selective activity against biofilms offers opportunities for potential applications, such as the titanium- tannin composite coating for implants developed by Shukla *et al*. (2015) (47), that allows sustained release of the tannin.

**FIG 3.**
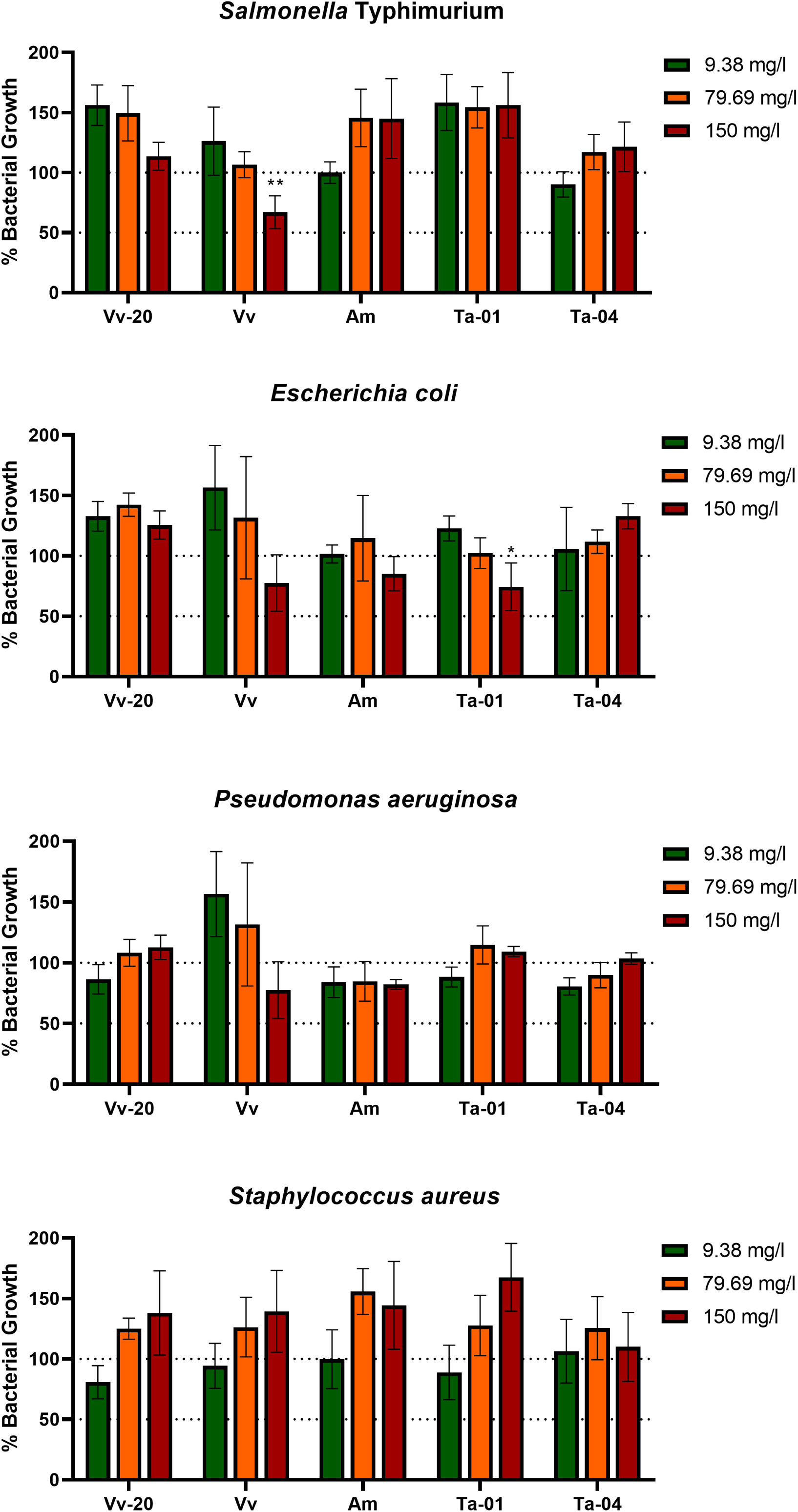
Planktonic growth (expressed as percentage in comparison to control) in the presence of 9.38, 79.69 and 150 mg/l of unmodified tannins. Asterixis indicate a significant difference from control, * = p < 0.05, ** = p < 0.01. The statistical differences were determined via ANOVA test with Tukey post-hoc analysis, with a p value of 0.05.

### Effect of chemical substitutions on the antibiofilm and antibacterial activity of the tannins

The aim of the modifications was to partially functionalize via ether linkages the phenolic OH-groups of each tannin molecule and to add tannin-alien functionalities at various levels to test the possibility to modulate the native activity of tannins towards biofilms and planktonic bacteria.

Fig. 4 shows the effect of derivatization on the antibiofilm and antibacterial activities. The differences in antimicrobial activity between the different modifications were assessed via the Tukey test for multiple comparisons, by comparing if there were differences in the maximum activity (i.e., the antibiofilm activity at the highest assayed concentration). If there were no differences in the maximum activity, the activity at lower concentrations was also evaluated, thus allowing to determine if a derivatization was able to obtain the same effect as the unmodified tannin, but at a lower concentration. As a general conclusion, it could be established that most of the chemical derivatizations, but especially positive charge introducing **C**_**3**_**NMe**_**3**_**Cl-0.1** reduce the antibiofilm activity of the tannins, while some of them can shift the activity spectrum towards preferential activity against Gram-positive or Gram-negative bacteria. Two derivatizations generally shifted the antibiofilm spectrum towards the Gram-negative group of bacteria: polymerizing **PEG**_**500**_**-0.05** and positive charge-introducing **C**_**3**_**NMe**_**3**_**Cl- 0.5**. Contrarily, derivatization with acidifying **C**_**3**_**COOH-0.1** and **C**_**3**_**COOH-0.5** in general decreased the antibiofilm activity against Gram-negative bacteria, while it retained activity against *S. aureus* for larger tannins **Am, Ta-01** and **Ta-04**, and increased activity against *S. aureus* for monomeric and low oligomeric **Vv-20** and **Vv**. The other derivatizations in general decreased the activity against *S. aureus*. However, this effect of derivatizations on the spectrum and potency was highly dependent on the specific tannin submitted to the derivatization.

**FIG 4.**
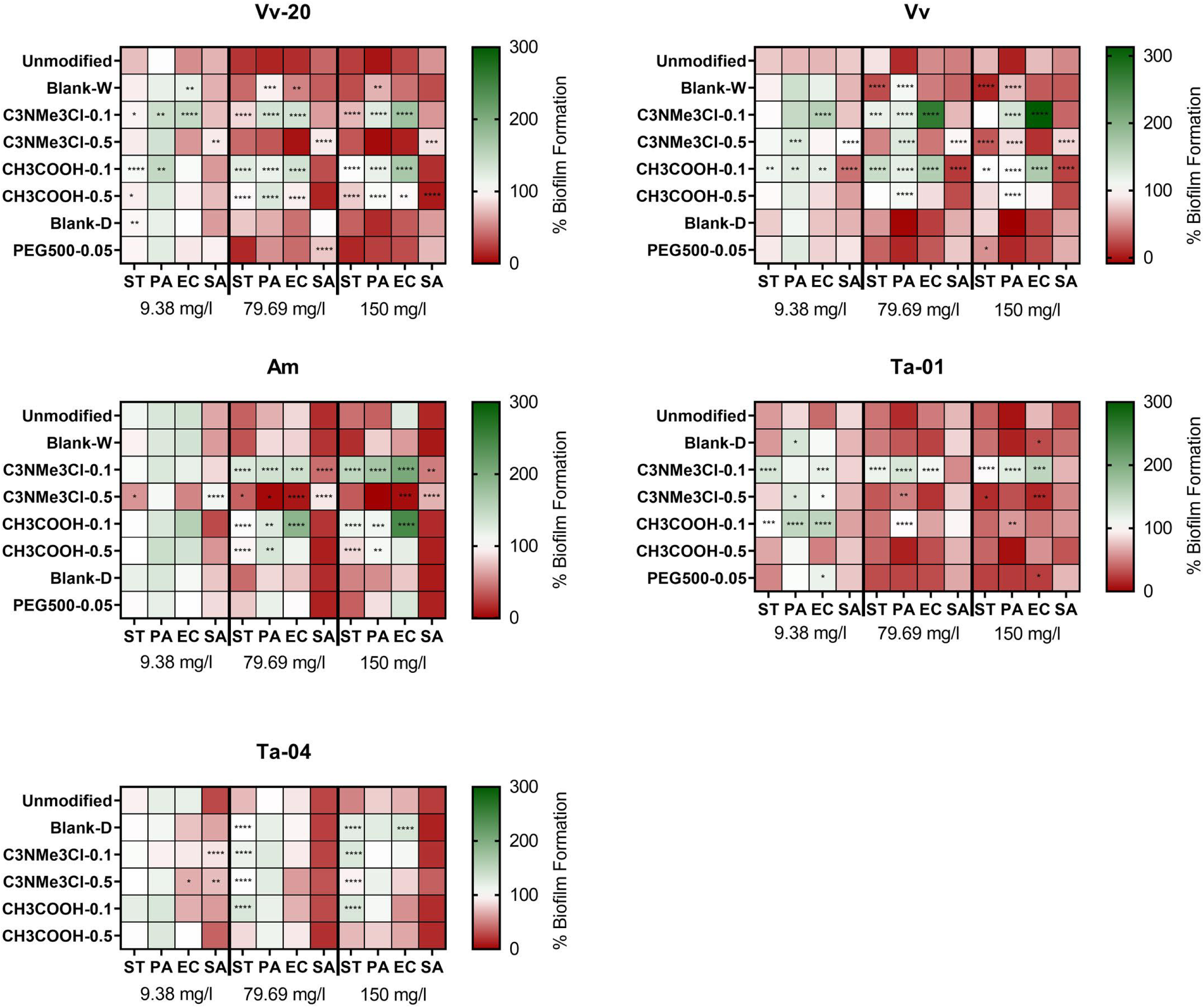
Effect of natural and chemically modified tannins on biofilm formation (expressed as percentage compared with positive control) on several bacterial species at 9.38 mg/l, 79.69 mg/l and 150 mg/l of tannin. The colors indicate the percentage of biofilm formation in presence of several concentrations of the assayed tannins compared to the untreated control. The asterisks indicate significant differences with the unmodified tannin, following ANOVA test with Tukey post-hoc analysis. *: p ≤ 0.05, **: p ≤ 0.01, ***: p ≤ 0.001, ****: p ≤ 0.0001.

In more detail, it can be seen that the blank reaction (both in water -**Blank-W**- and in dimethylformamide -**Blank-D**-) already modified the antibiofilm effect of the assayed tannins, a phenomenon that could be attributed to removal of impurities that affect the antibiofilm effect of the tannins. It can be seen that **Blank-W** conditions increased the maximum antibiofilm effect of **Vv** against *Salmonella* Typhimurium, but decreased the maximum antibiofilm effect of **Vv-20** and **Vv** against *P. aeruginosa* and decreases the antibiofilm effect of **Vv-20** against *E. coli* at 79.69 mg/l without affecting its maximum antibiofilm effect. It can also be seen that **Blank-D** conditions only affected the antibiofilm activity of **Ta-04**, by reducing its antibiofilm effect against Gram-negative bacteria.

Regarding actual substitutions, it can be seen that the effect of a derivatization with positive charge introducing ammonium groups, i.e., **C**_**3**_**NMe**_**3**_**Cl**, was different depending on the equivalents of chemical substitution.

On the one hand, **C**_**3**_**NMe**_**3**_**Cl-0.5**, the high equivalent derivatization, decreased the maximum antibiofilm effect of condensed tannins against *S. aureus* but did not affect the anti-staphylococcal effect of hydrolysable tannins. However, **C**_**3**_**NMe**_**3**_**Cl-0.5** derivatization had a tannin-dependent effect against Gram-negative bacteria. It did not affect the antibiofilm effect of monomeric **Vv-20** against any Gram-negative bacteria but increased the maximum antibiofilm effect of low oligomeric **Vv** against *Salmonella* Typhimurium while decreasing the maximum antibiofilm effect of **Vv** against *P. aeruginosa*. It increased the effect of high oligomeric **Am** against all the Gram-negative bacteria at 79.69 mg/l without significantly changing the maximum effect compared to the unmodified tannin. It increased the maximum effect of tannic acid **Ta-01** against *E. coli* and *Salmonella* Typhimurium but decreased the effect against *P. aeruginosa* at 79.69 mg/l and it significantly decreased the maximum antibiofilm effect of galloquinic acid derivative **Ta-04** against *Salmonella* Typhimurium. On the other hand, **C**_**3**_**NMe**_**3**_**Cl-0.1**, the low equivalent derivatization, reduced the maximum effect of all tannins (with the exception of the galloquinic acid derivative **Ta-04**, which was not affected) against Gram-negative bacteria without affecting the antibiofilm effect against *S. aureus*.

Contrary to **C**_**3**_**NMe**_**3**_**Cl**, there were no big differences between different levels of derivatizations with acidifying motif **CH**_**3**_**COOH**, i.e., **CH**_**3**_**COOH-0.1** and **CH**_**3**_**COOH-0.5**, in terms of their impact on the activities compared to the unmodified tannins. For *S. aureus*, neither derivatization level affected the antibiofilm effect exerted by larger tannins **Am, Ta-01**, and **Ta-04**, comparable to the **C**_**3**_**NMe**_**3**_**Cl-0.1** derivatization, while **CH**_**3**_**COOH-0.1** increased the maximum effect of low oligomeric **Vv. CH**_**3**_**COOH-0.5** had a similar effect on essentially monomeric **Vv-20**. For the Gram-negative bacteria, both derivatizations equally decreased the maximum antibiofilm effect of condensed tannins **Vv-20** and **Am**, as well as for hydrolysable tannin **Ta-04** against *S*. Typhimurium, but only **CH**_**3**_**COOH-0.1** significantly decreased the maximum antibiofilm effect against *S*. Typhimurium of **Vv**. While both derivatization levels drastically decreased the maximum antibiofilm effect of condensed tannins against *P. aeruginosa*, only **CH**_**3**_**COOH-0.1** decreased the maximum antibiofilm effect of tannic acid **Ta-01**. Neither derivatization level significantly affected the antibiofilm effect of hydrolysable **Ta-04**. This effect was similar regarding *E. coli* because neither derivatization level affected the maximum antibiofilm effect of hydrolysable tannins, while **CH**_**3**_**COOH-0.1** decreased the maximum antibiofilm effect of all condensed tannins. **CH**_**3**_**COOH-0.5** only significantly decreased the maximum antibiofilm effect of **Vv-20** against *E. coli*.

Regarding derivatization with polymerizing **PEG500**, it can be seen that **PEG500-0.05** in general did not affect the antibiofilm effect of the assayed tannins. The only exceptions are the increase in the maximum antibiofilm effect of low oligomeric **Vv** against *Salmonella* Typhimurium and the decrease in the antibiofilm effect of monomeric **Vv-20** against *S. aureus* at 79.69 mg/l without changing the maximum antibiofilm effect. It has to be taken in account that, due to technical issues, it was not possible to analyze the effect of polymerizing **PEG500-0.05** derivatization on **Ta-04**. Polymerizing **PEG500-0.05** derivatization proofed difficult to analyze in terms of loading for **Ta-04** (41).

No literature data are yet available that would describe the effects of chemical modifications of tannins on their antibiofilm, or more generally antibacterial activity. With the aim of elucidating a hypothesis about the reason behind the effect of the derivatizations on our assayed tannins in comparison to the parent tannins and the blanks, we decided to look into other non-tannin organic compounds.

One of these examples targets the effect of the high equivalent derivatization **C**_**3**_**NMe**_**3**_**Cl-0.5** in shifting the anti-biofilm spectrum towards Gram-negative bacteria, which may be associated with addition of positive charges in the form of ammonium groups to the tannin molecules. This allows for comparison of our results with data obtained by Dalcin *et al*. (2017) (48), who discovered that nanoencapsulation of dihydromyricetin within the polycationic polymer Eudragit RS 100® not only increased its antibiofilm activity against *P. aeruginosa*, but that the polymer itself had antibiofilm activity. This finding goes in accordance with a previous publication of Campanac *et al*. (2002) (49) which states that cationic quaternary ammonium compounds (QAC) were more effective against *P. aeruginosa* than *S. aureus* biofilm. Also, Gao *et al*. (2019) (50) showed a decreased biofilm formation in *E. coli* and *S. aureus* using positively charged nanoaggregates based on zwitterionic pillar-[5]arene, requiring a ten times lower concentration of nanoaggregate to decrease biofilm formation in *E. coli* compared to *S. aureus*. These results suggest that equipping the tannins with positively charge-introducing ammonium groups may give preferential action against Gram-negative bacteria. However, tannins should be derivatized with enough positively charge-introducing ammonium groups to obtain this shift towards inhibition of Gram-negative bacteria since the low equivalent derivatization **C**_**3**_**NMe**_**3**_**Cl-0.1** appeared to lower the activity against Gram-negative bacteria.

Regarding the effect of acidifying **CH**_**3**_**COOH** derivatizations, there is a precedent of the effect of several substitutions on the antibiofilm effect of anthraquinones against methicillin-resistant *S. aureus* (MRSA) (51). This study shows that a carboxyl group at position 2 of the anthraquinone molecule increases both the antibiofilm and the antimicrobial activity, which is partially in agreement with our data that shows that **CH**_**3**_**COOH** derivatizations increase the antibiofilm activity of low oligomeric **Vv-20** and essentially monomeric **Vv** without affecting the bacterial growth. Relatedly, Warraich *et al*. (2020) (52) found that the acidic D-amino acids D-aspartic acid (D-Asp) and D-glutamic acid (D-Glu) were effective in dispersing and inhibiting biofilm formation in *S. aureus*, and they attributed this effect to the negative charges introduced by carboxyl groups of the molecules under growth conditions.

Finally, regarding polymerizing **PEG** derivatizations, there are several studies about the potential of PEG cross-linked hydrogels for wound healing because of their antimicrobial, pro-angiogenesis and pro-epithelization capabilities (53), but there is no indication regarding the effect of an introduction of a PEG-motif on the antibiofilm capability of an organic compound.

### Biofilm specificity of the antibacterial effect of the tannins

In the last years, there has been increasing research on non-lethal antimicrobial targets against several bacterial species, from virulence factors to biofilm formation, including inhibition of regulatory mechanisms such as quorum sensing and production of public goods (54–57). The rationale behind this research is the assumption that if bacterial viability is not affected, the selective pressure will be lower and thus the risk for emergence of antimicrobial resistance will be lower too (35, 58, 59). However, there is still discussion about the effectiveness of this “resistant-proof” approach (60).

The heatmap of Fig. 5 shows the anti-planktonic activity of the tannins at the assayed concentrations. We defined that the antibiofilm activity of a particular tannin was biofilm specific in case the tannin did not exhibit significant anti-planktonic effect at that concentration (61, 62). Most of the tannins with antibiofilm activity did not exhibit anti-planktonic effects, indicating that they are biofilm specific. This is particularly true for unmodified tannins, of which only **Vv** and **Ta-01** exhibited anti-planktonic activity against *Salmonella* Typhimurium and *E. coli*, respectively. Also, unmodified tannins re-isolated from blank reactions, i.e., **Blank-W** or **Blank-D**, with the exception of **Blank-D** against *Salmonella* Typhimurium, did not reduce the planktonic growth of the assayed bacteria.

**FIG 5.**
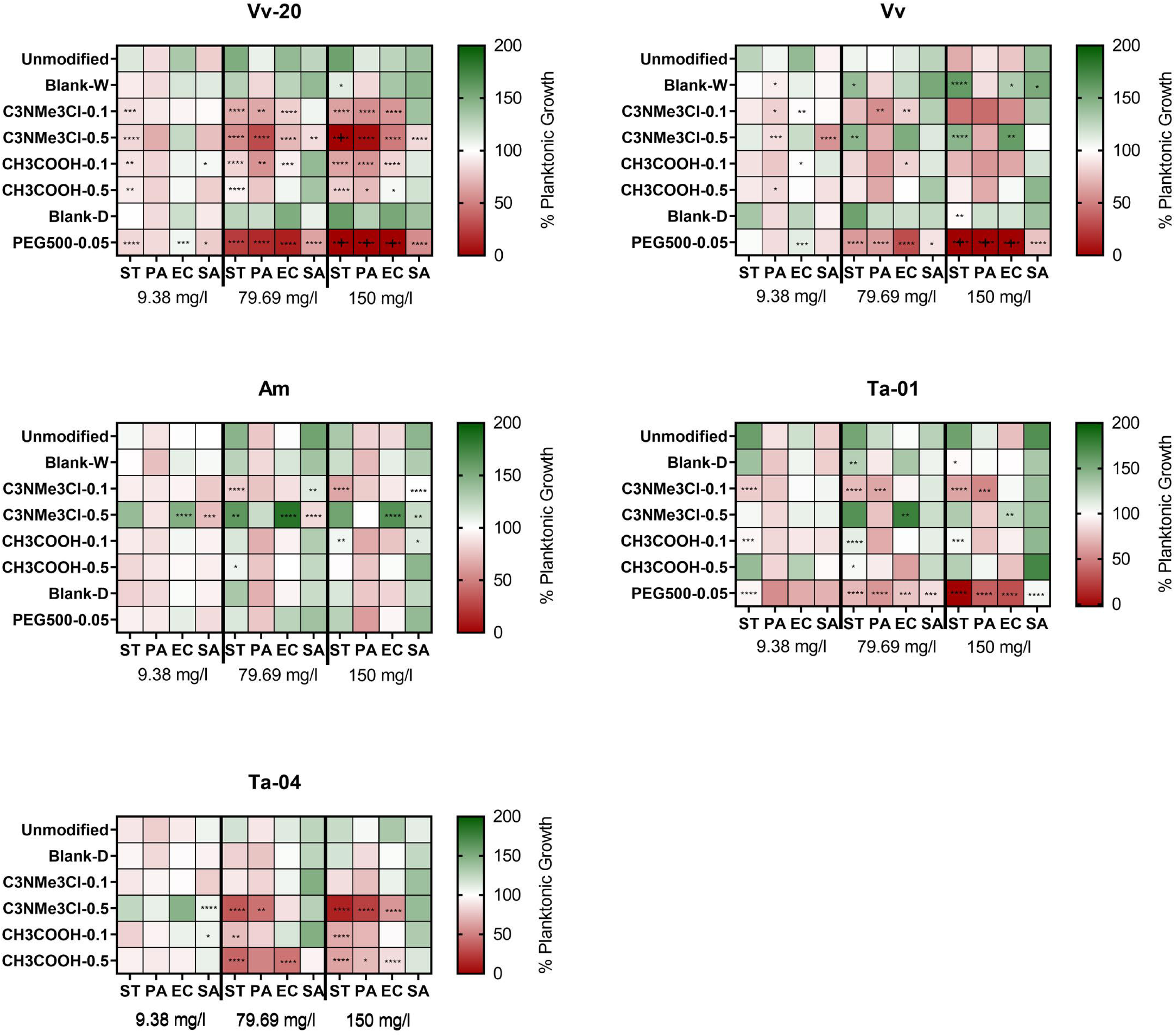
Effect of natural and chemically modified tannins on planktonic growth (expressed as percentage compared with control) on several bacterial species at 9.38 mg/l, 79.69 mg/l and 150 mg/l of tannin. The crosses (**+**) for **PEG**_**500**_**-0.05** derivatization on **Vv-20** and **Vv** indicate values below zero, which is an effect of potential overcorrection of the raw values by the negative control. The colors indicate the percentage of planktonic growth in presence of several concentrations of the assayed tannins compared to the untreated control. The asterisks indicate significant differences with the unmodified tannin, following ANOVA test with Tukey post-hoc analysis. *: p ≤ 0.05, **: p ≤ 0.01, ***: p ≤ 0.001, ****: p ≤ 0.0001.

Regarding the effect of derivatizations on the biofilm specificity of the tannins, this was both derivatization and species dependent. Derivatization with positive charge introducing **C**_**3**_**NMe**_**3**_**Cl-0.1** not only generally decreased the antibiofilm activity of the assayed tannins, but also increased the anti-planktonic activity specifically against Gram-negative bacteria. Antibiofilm specificity of tannins derivatized with **C**_**3**_**NMe**_**3**_**Cl-0.5** was highly dependent on both the assayed bacteria and the derivatized tannin: the antibiofilm effect was non-specific when the derivatization was applied to **Vv-20**, but it was biofilm-specific for **Vv, Am** and **Ta-01**. Also, tannins derivatized with **C**_**3**_**NMe**_**3**_**Cl-0.5** most strongly affected the growth of *P. aeruginosa* except for **Am-C**_**3**_**NM3e**_**3**_**Cl-0.5**, which did not affect the growth of *P. aeruginosa* at any concentration. Only a derivatization of **Ta-04**, and here especially with **C**_**3**_**NMe**_**3**_**Cl- 0.5**, led to a dose-dependent anti-planktonic effect, thus exhibiting non-specific antibiofilm activity at higher concentrations. With respect to a functionalization with the acidifying element **CH**_**3**_**COOH** at various concentrations, it can be stated that these in this study did not significantly change the anti-planktonic behavior of the derivatized **Vv, Am** and **Ta-01** against the assayed bacteria, but increased the antibacterial effect of **Vv-20** and **Ta-04** against planktonic bacteria. However, one notable exception are the tannins that were modified with crosslinking **PEG500-0.05**, whose antibiofilm activity against Gram negative bacteria was highly correlated with the ability to inhibit the planktonic growth (Fig. 5).

As a general summary, tannin derivatization did not affect biofilm specificity against *S. aureus* but affected the biofilm specificity against Gram-negative bacteria in a tannin-specific and derivatization specific manner. More importantly, there was no correlation in the assayed tannins between the degree of inhibition of planktonic growth and the degree of antibiofilm effect, a situation that goes in accordance with some previous reports (34, 63, 64).

### Relation between antibiofilm effect and phenolic hydroxyl content of the tannins

One of the potential consequences of the chemical derivatization of the tannins are changes in the content of free phenolic hydroxyl groups present in the chemical structure of the tannins, since functionalization occurs at these hydroxyl groups. This is potentially important, because in previous literature it has been described that the biological activity of polyphenols could be mediated by their phenolic hydroxyl groups (65–71). Particularly regarding hydrolysable tannins, Taguri *et al*. (2004) linked the degree of antibacterial activity to the presence of galloyl groups (6). Because of this, we aimed to determine if there was a significant impact of the derivatizations on the phenolic hydroxyl (OH) content of the tannins, and if the phenolic OH content affected the antibiofilm.

In a first approach, we studied the impact of derivatizations on the phenolic OH content of the tannins. As can be seen in Table 1, derivatizations with **C**_**3**_**NMe**_**3**_**Cl-0.1** decreased the phenolic OH content of the unmodified tannin. This is important, since derivatization with **C**_**3**_**NMe**_**3**_**Cl-0.1** significantly decreased the antibiofilm effect of tannins, mostly against Gram negative bacteria. In a similar trend, derivatizations with **CH**_**3**_**COOH** tended to decrease the phenolic content of condensed tannins: derivatization with **CH**_**3**_**COOH** decreased the antibiofilm effect against Gram negative bacteria of condensed tannins, but, interestingly, not of still larger hydrolysable tannins.

**TABLE 1.** Phenolic OH content (in mmol phenolic OH / g materials) of unmodified and derivatized tannins. The values were obtained using ^31P^NMR.

| Tannin | Phenolic OH content |  |  |  |  |  |  |  |
| --- | --- | --- | --- | --- | --- | --- | --- | --- |
|  | Unmodified | Blank-W | Blank-D | C<sub>3</sub>NMe<sub>3</sub>Cl-0.1 | C<sub>3</sub>NMe<sub>3</sub>Cl-0.5 | CH<sub>3</sub>COOH-0.1 | CH<sub>3</sub>COOH-0.5 | PEG<sub>500</sub>-0.05 |
| **Vv-20** | 10.37 | 7.85 | 8.79 | 4.35 | N.d.<sup>a</sup> | 6.19 | 6.15 | 7.03 |
| **Vv** | 5.38 | 6.83 | 11.70 | 3.26 | 5.21 | 1.51 | 5.09 | 7.11 |
| **Am** | 8.35 | 7.64 | 10.27 | 0.45 | N.d.<sup>a</sup> | 5.76 | 3.76 | 8.03 |
| **Ta-01** | 12.56 | ▪<sup>b</sup> | 10.99 | 1.55 | 7.22 | 10.05 | 11.61 | 8.87 |
| **Ta-04** | 10.49 | ▪<sup>b</sup> | 6.89 | 6.00 | N.d.<sup>a</sup> | 9.89 | 9.00 | 6.77 |

<sup>a</sup> N.d.: Not possible to calculate phenolic OH content due to solubility issues.
<sup>b</sup> ▪ Blank-W reaction was not performed with Ta-01 and Ta-04 (hydrolysable tannins) because, for those tannins, it was not possible to perform a blank reaction in water.

In a second approach, we hence studied potential correlations between the phenolic OH content and the antibiofilm activity and we observed a weak correlation between the phenolic OH content and the antibiofilm activity against Gram negative bacteria and, to a lesser extent, against *S. aureus* (see Fig. S3 in ‘Supplementary Material’). These data, combined with the previously mentioned effect of the different derivatizations on the phenolic OH content of the tannins, suggest that the phenolic OH content of the tannins is more important for the antibiofilm effect against Gram negative bacteria than against *S. aureus*. However, we did not observe a correlation between the antibacterial activity against planktonic bacteria and the phenolic OH content of the assayed tannins (see Fig. S4 in ‘Supplementary Material’), which is in accordance with the study Kim *et al*. (2020) (72), who did not find a significant correlation between the total phenolic content of several plant extracts from Chinese traditional medicine and the antimicrobial activity against *S. aureus*. However, this contradicts a previous study from Vattem *et al*. (2004) (73), who found a linear correlation between the phenolic content of cranberry pomace and the antimicrobial activity against *Listeria monocytogenes, Vibrio parahaemolyticus* and *E. coli*. It is important to note that these studies are focused on planktonic bacteria, since there are no previous studies pointing to the effect of the hydroxyl or phenolic content of bioactive compounds on biofilm inhibition and dispersion.

### Conclusions

Our work provides a clear understanding on which chemical modifications can be made to enhance the activity level or change the activity spectrum of natural tannins, and which chemical modifications are inconvenient for increasing their antibiofilm activity. This is one of the few studies that uses a systematic and statistical analysis to correlate specific chemical characteristics of modified tannins with their antibiofilm activity (38, 74–78).

From our work, it can be concluded that tannins not only have good activity against biofilms of different bacterial species, but that this antibiofilm activity is in most cases also biofilm specific. More important, we could identify that modifying the tannins with **C**_**3**_**NMe**_**3**_**Cl-0.5** and **PEG500-0.05** can increase the antibiofilm activity against Gram-negative bacteria, although this often coincides with a decrease in activity against Gram-positive bacteria. Modifying the tannins with **C**_**3**_**NMe**_**3**_**Cl-0.1, CH**_**3**_**COOH- 0.1** and **CH**_**3**_**COOH-0.5** generally decreases the effect against Gram-negatives, without affecting the activity against *S. aureus*. We can thus modulate the spectrum and the antibiofilm potency of tannins by the applied chemical modifications.

We could identify a weak correlation between the antibiofilm effect and the content of phenolic hydroxyl groups for Gram-negative bacteria and, to a lesser extent, for *S. aureus*. However, exploring the mode of action against other bacteria is a necessary and interesting avenue to explore further based on the initial insights generated in this work, pointing to a more complex interplay between functionalization, type of tannin in the sense of exposed galloyl units, and tannin size. A continued exploration of the possible mechanisms of actions of these compounds and the possible modifications that can be made to enhance their effect is necessary to better optimize the antibiofilm potential of the tannins.

## Acknowledgements

We especially like to thank David De Coster for his technical support during the experimental work.

## Transparency declarations

The authors declare no conflict of interest, and that all the published information is truthful to the results of our experimental procedures.

